# Particulate Matter fractions and kitchen characteristics in Sri Lankan households using solid fuel and LPG

**DOI:** 10.1101/461665

**Authors:** Sumal Nandasena, Rajitha Wickremasinghe, Anuradhani Kasturiratne, Udaya Wimalasiri, Meghan Tipre, Rodney Larson, Emily Levitan, Nalini Sathiakumar

## Abstract

**Introduction:** Use of solid fuel for cooking is a major source of household air pollution in developing countries. Of the many pollutants emitted during solid fuel combustion, Particulate Matter (PM)is considered to be one of the most hazardous pollutants. We monitored PM fractions emitted during solid fuel and Liquefied Petroleum Gas(LPG) combustion in kitchens of Sri Lankan households.

**Methods:** Households of children in a longitudinal study in Ragama, Sri Lanka was the study population. At the age of 36 months of children, a sample of households were visited and different aerodynamic diameters of PM (PM_1,_ PM_2.5_, PM_10_) were monitored during the main cooking session for 3 hours. Basic characteristics of kitchen (e.g., availability of chimney, functionality of chimney, etc.) were assessed by a questionnaire. Cooking energy, other sources of household air pollution, size of open spaces in the kitchen (e.g., windows), etc. were assessed at the time of PM monitoring.

**Results:** Questionnaire was administered for mothers in 426 households. Out of them, 245 (57.5%)and 116 (27.2%) households used LPG and wood as the primary cooking fuel respectively. During the cooking period, PM_2.5_ concentrations of households uses only wood fuel and cook inside the main housing building were 344.1 μg/m^3^(Inter Quartile Range(IQR) = 173.2-878.0μg/m^3^), 88.7 μg/m^3^(54.8- 179.2 μg/m^3^); 91.7 μg/m^3^ (56.0- 184.9 μg/m^3^) and 115.1 μg/m^3^(83.4 - 247.9 μg/m^3^) in kitchen, sleeping room, living room and immediate outdoor respectively. Immediate outdoor PM_2.5_ concentrations in wood burning households was higher among households not having chimney (n = 8)compare to those having a chimney (n = 8) (245.9μg/m^3^ (IQR = 72.5 – 641.7μg/m^3^)) VS. (105.7μg/m^3^ (83.4– 195.8μg/m^3^)).Fuel type and stove type, availability of a chimney and their functional status, ratio between open space and total space of kitchen, PM_2.5_ concentration at the non-cooking time (i.e., baseline PM_2.5_concentration) were the determinants of PM_2.5_ in wood using kitchens during cooking period.

**Conclusions:** PM concentrations were higher in kitchen and other microenvironments of the households use wood for cooking as compared to LPG use for cooking. Immediate outdoor PM concentration was higher than the sleeping and living room PM concentrations. Several factors determine the PM_2.5_concentrationsduring the cooking including the fuel type.

## Introduction

Solid fuel use for cooking is a major source of household air pollution, and is considered to be a leading cause of adverse health effects [1]. Household air pollution is predominantly reported with solid fuel use for cooking [2]. Solid fuel combustion is linked to an array of adverse health effects [4-6]. It is estimated that 4.3 million premature deaths are attributable to household air pollution [1]. About 66% of Sri Lankan households use solid fuel as the primary source of cooking fuel. The use of solid fuel for cooking has not declined markedly over the years [2]. The most commonly used solid fuel in Sri Lanka is wood (2). In Sri Lanka, solid fuel use was least in the urban sector (25% of households) while the percentage was highest in the plantation sector (80%) [4].

The health effects of air pollutants emitted from solid fuel cook stoves depend on the type of solid fuel used, the concentration of the pollutant, duration and frequency of exposure and toxicity associated with the pollutants. Further, concentration of pollutants depends on kitchen characteristics such as ventilation, presence of a chimney and stove design [5]. Commonly used stove types in Sri Lanka include traditional stoves as well as improved cook stoves (i.e., “Anagi” stove). “Traditional stoves” which have been used in households for many years are mostly user built with locally available materials such as the “three stone stove”. “Anagi stoves” are light weight clay products that could accommodate one or two cooking utensils at a time and are commercially available (Figure 1). Previous studies have reported that the use of improved cook stoves and presence of a chimney reduces personal exposure to PM [5].

**Figure 1:**
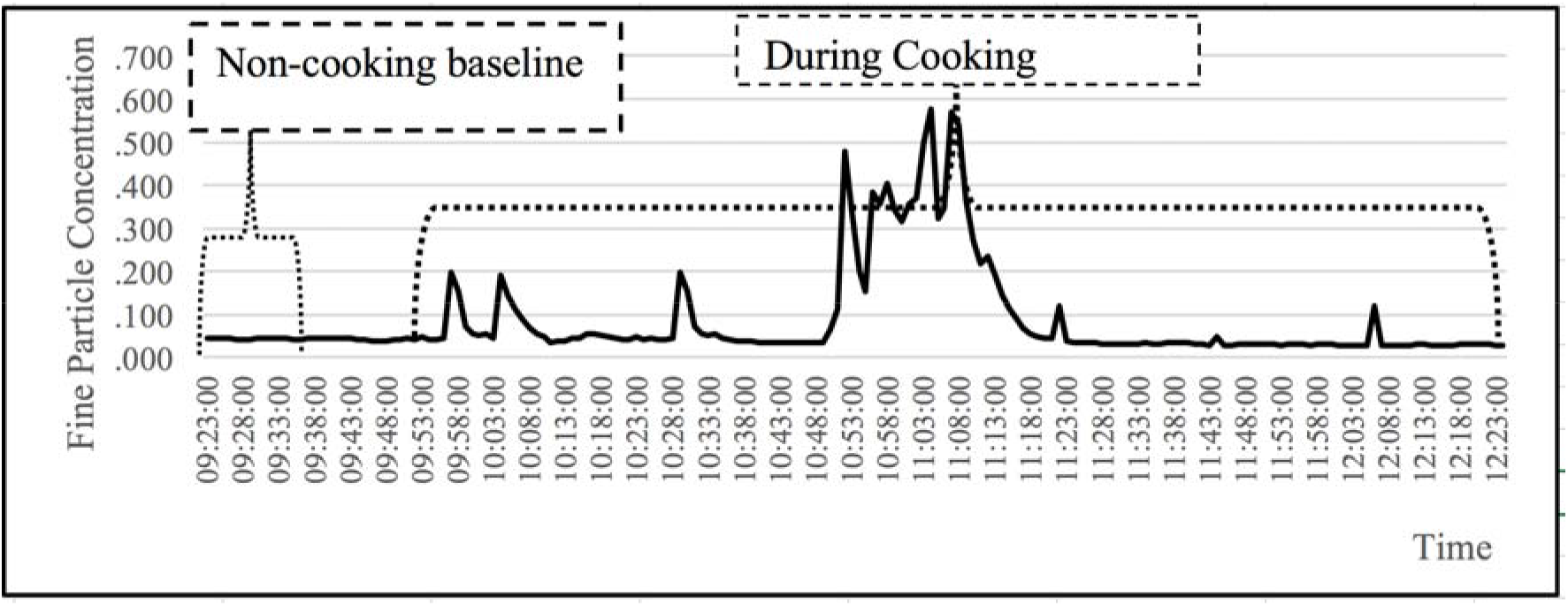
Estimation of PM concentrations at baseline and during cooking.

PM, a mixture of solid and liquid particles suspended in the air, is one of the most hazardous pollutants emitted during combustion of solid fuel [7]. PM is directly emitted into the air or forms in the atmosphere from gaseous precursors [8]. PM produces adverse health effects via different mechanisms. For example, PM increases susceptibility to infection through alveolar macrophage-driven inflammation, alterations in alveolar macrophage phagocytosis and up-regulation of receptors involved in pathogen invasion [9].

PM is commonly described by the size fraction of its aerodynamic diameter. PM_10_ refers to PM with an aerodynamic diameter < 10 μm and PM_2.5_ refers to PM with an aerodynamic diameter <2.5 μm. PM is quantified by its mass concentration in the air (μg/m^3^)[8]. PM with smaller aerodynamic diameters are more hazardous as they are able to penetrate deep into the lungs [7].

In this study, we describe PM_1,_ PM_2.5_ and PM_10_ concentrations during cooking using fire wood and LPG in different microenvironments (i.e., kitchen, living room and sleeping room) and the immediate outdoors of Sri Lankan households. Further, we describe important determinants of PM_2.5_ concentrations in kitchens.

## Methodology

This study is part of a larger prospective study examining the effect of biomass fuel use on birth outcomes and development of children under five years. This study describes the exposure related information ascertained by an interviewer administered questionnaire and monitoring of PM in households.

### Study Setting

The study was conducted in the Ragama Medical Officer of Health area, an area with a multi-ethnic population having semi-urban to rural characteristics in the Gampaha district of Sri Lanka. The Gampaha district is the second most populous district of the country with population density of 1589/ km^2^. Study households were located over a 25km^2^ area of the Gampaha District.

### Data collection

Appointments for home visits on a convenient day were made beforehand in consultation with the mother of the child followed up in the larger study. An interviewer administered questionnaire was administered to mothers of 426 households after obtaining informed written consent. The questionnaire included extensive details on household characteristics and practices related to household air pollution exposure, socio-demographic data, etc. Information on type of cooking fuel used (primary and secondary), location of the kitchen (i.e., indoors or outdoors), size of the kitchen, access to ventilation of the kitchen was estimated as the area of open spaces in the kitchen (i.e., area of doors, windows and other openings), stove type (traditional vs. improved stoves), etc. was also collected.

PM_1.0,_ PM_2.5_, PM_10,_ were monitored in a subsample of houses during preparation of lunch, the main meal of the day, on a day convenient to the householders. Monitors were fixed in the kitchen, the living room and the sleeping room of the household, and in the immediate outdoors of the kitchen. A standard monitoring protocol was used to position the monitors such that the air inlets of the monitors were fixed at 145cm above the floor level (i.e., breathing zone of an adult). PM was monitored using DustTrak™ DRX aerosol monitor 8533 which monitors PM fractions simultaneously. Monitoring was started approximately 20 - 30 minutes prior to the commencement of the cooking session.

In addition to the baseline questionnaire, a check list was used at the time of air quality monitoring to note down specific conditions present during PM monitoring. The type(s) of fuel used during the monitoring session as well as the presence of other sources of household air pollution such as presence of smokers in the kitchen or surrounding areas were recorded. The exact time of starting the cooking session was noted enabling the calculation of baseline PM concentrations.

## Statistical Analysis

As air quality data were skewed, the median and the interquartile range were used to describe data. The minute to minute logged data was used to calculate (1) PM concentration prior to the cooking session (baseline) and (2) PM concentration during the cooking period (cooking)as shown in figure 1. Baseline PM concentration was taken during the first 10 minutes of monitoring. Data from the end of baseline to the start of the cooking period were not taken as pollutant levels may change when the cook stove is being cleaned and set up. Information on the start of the cooking time was obtained from the check list. Then, PM concentrations were monitored from the start of cooking to the end of the monitoring period. Households were monitored for approximately 2 1/2 hours. PM concentrations at baseline and during cooking were compared using the Mann-Whitney U Statistic). PM concentrations were log transformed and a multiple linear regression model fitted.

## Ethical Approval

Ethical approval was obtained from the Ethics Review Committee of the Faculty of Medicine, University of Kelaniya, Sri Lanka and the School of Public Health of the University of Alabama at Birmingham. Informed written consent was obtained from the mother at the first point of contact after explaining the study objectives and procedures. Verbal consent was obtained prior to the home visits and PM monitoring. A convenient date for the family was scheduled for the home visit. Permission was obtained from the local health authorities.

Mothers residing in houses with high levels of PM were advised on how to mitigate the adverse effects of indoor air pollution.

## Results

Kitchen characteristics of the entire study population (n = 426 households) are given in table 1; 245 (57.5%) households used LPG as the primary cooking energy (i.e., cooking fuel use > 75%) while 116 (27.2%) households used wood as the primary cooking energy (i.e., cooking fuel use > 75%). Considerable number of households include more than one energy for cooking. Less than 25% of energy source for a cooking sessions was electricity (for rice cookers, kettles, etc.) in 44% (n = 188) of households. About 47% (n=54) of households that predominantly used wood as the cooking fuel were using multiple pot improved stoves. The majority of the households had a separate room (n = 368, 86.4%)(kitchen) for cooking in the main building. Over 50% of households that cook indoors had a chimney.

**Table 1:** Kitchen characteristics of the study population (information from baseline questionnaire)

| **Kitchen Characteristics (n=426)** | **n** | **%** |
| --- | --- | --- |
| **Place of cooking** |  |  |
| Same room used for sleeping: un-partitioned | 8 | 1.9 |
| Same room used for sleeping: partitioned | 3 | 0.7 |
| Separate room in the main building (kitchen) | 368 | 86.4 |
| Separate building (kitchen not in main house) | 4 | 0.9 |
| Outdoors or in a temporary hut | 43 | 10.1 |
| **Chimney(n=383)\*** |  |  |
| Yes | 199 | 52.0 |
| No | 184 | 48.0 |
| **Condition of chimney (n=199)** |  |  |
| Very good condition | 56 | 28.14 |
| Good condition | 78 | 39.19 |
| Fair condition | 46 | 23.11 |
| Poor condition | 19 | 9.5 |
| **Primary cooking fuel ($\geq 75\%$ of the time)** |  |  |
| Electricity | 1 | 0.2 |
| LPG | 245 | 57.5 |
| Kerosene | 7 | 1.6 |
| Wood | 116 | 27.2 |
| Dry vegetation | 2 | 0.4 |
| **Secondary cooking fuel (<25% of the time)\*\*** |  |  |
| Electricity | 188 | 44.1 |
| LPG | 46 | 10.8 |
| Kerosene | 3 | 0.7 |
| Wood | 50 | 11.7 |
| Dry vegetation | 182 | 42.7 |
| **Lighting** |  |  |
| Electricity | 425 | 99.8 |
| Kerosene | 1 | 0.2 |

| **Primary cook stove type(n = 116) \*\*\*** |  |  |
| --- | --- | --- |
| Open fire | 11 | 9.4 |
| Protected fire | 27 | 23.3 |
| Improved single pot stove | 24 | 20.7 |
| Improved multiple pot stove | 54 | 46.6 |

\* Households that cook indoors
\*\* Households predominantly using wood for cooking
\*\*\*Households using wood as the primary cooking energy **>75% of the time**

PM monitoring was done in a subsample of households. PM_2.5_concentrations (at baseline and during cooking) were monitored in microenvironments within the households and in the immediate outdoors of households (n =146)(table 2). Of these, 48 households used only wood (i.e., 100% wood use), 64 households used only LPG (i.e., 100% LPG use), 21 households used wood for between 50-75% of the time and 13 households used wood for more than 75% of the time but less than 100% of the time. The highest PM_2.5_ concentration during cooking was reported in kitchens in households using wood 100% of the time (median=344.1 (IQR = 173.2-878.0 μg/m^3^)). There was no difference in the baseline PM_2.5_ concentrations in any of the microenvironments between households using 100% wood and 100% LPG. However, there were significant differences in PM concentrations during the cooking period in all microenvironments between households using 100% wood and 100% LPG.

**Table 2:** PM concentrations in different microenvironments of households and immediate outdoors.

|  |  | PM concentrations at baseline |  |  | PM concentrations during cooking |  |  |
| --- | --- | --- | --- | --- | --- | --- | --- |
| Cooking fuel type and location of monitoring | n | PM<sub>1</sub><br>median (IQR)<br>µg/m<sup>3</sup> | PM<sub>2.5</sub><br>median (IQR)<br>µg/m<sup>3</sup> | PM<sub>10</sub><br>median (IQR)<br>µg/m<sup>3</sup> | PM<sub>1</sub><br>median (IQR)<br>µg/m<sup>3</sup> | PM<sub>2.5</sub><br>median (IQR)<br>µg/m<sup>3</sup> | PM<sub>10</sub><br>median (IQR)<br>µg/m<sup>3</sup> |
| **Kitchen** |  |  |  |  |  |  |  |
| 100% wood use | 48 | 55.3(36.5-105.7) | 60.5 (40.1-114.21) | 75.2 (47.4-122.3) | 337.4 (166.4-877.1) | 344.1 (173.2-878.0) | 348.1 (185.21-879.3) |
| > 75% to <100% wood use | 13 | 80.13(30.64-211.1) | 80.97 (32.47-214.65) | 82.03 (36.23-217.25) | 310.2 (197.6-535.6) | 314.5 (203.2-565.5) | 328.1 (218.5-586.3) |
| 50% to 75% wood use | 21 | 51.8 (32.3-105.3) | 59.7 (37.3.1-108.8) | 77.2 (44.8-123.0) | 225.7 (106.8 – 756.7) | 233.6 (121.9-769.7) | 248.3 (136.6 – 788.7) |
| 100% LPG use | 64 | 50.2 (29.1-77.3) | 52.5 (33.4-79.2) | 58.3 (43.3-85.9) | 51.8 (37.8-98.9) | 53.6 (39.7-78.0) | 69.5 (47.2 -103.2) |
| p-value<sup>1</sup> |  | 0.34 | 0.34 | 0.34 | <0.001 | <0.001 | <0.001 |
| **Living Room<sup>2</sup>** |  |  |  |  |  |  |  |
| 100% wood use | 41 | 57.2(31.1-86.1) | 61.2(33.3-88.3) | 64.4(36.1-91.7) | 90.8(54.6-182.5) | 91.7(56.0-184.9) | 93.6(59.3-188.4) |
| > 75% to <100% wood use | 10 | 47.7(23.3-106.3) | 53.3(25.2-107.1) | 65.0(39.5-109.0) | 83.2(33.2-146.7) | 83.5(36.4-151.0) | 84.6(45.0-153.5) |
| 50% to 75% wood use | 19 | 44.7(29.4-76.3) | 48.2(30.4-77.2) | 52.2(32.3-80.2) | 69.4 (39.1- 126.3) | 71.3 (39.2 – 127.1) | 71.8 (39.6 – 138.1) |
| 100% LPG use | 56 | 53.7(30.5-89.9) | 56.8(33.1-91.3) | 61.4(37.4-93.8) | 48.3(32.1-80.7) | 49.9(33.9-81.3) | 51.4(38.0-82.3) |
| p-value<sup>1</sup> |  | 0.931 | 0.635 | 0.931 | <0.001 | <0.001 | <0.001 |
| **Sleeping room<sup>2</sup>** |  |  |  |  |  |  |  |
| 100% wood use | 47 | 55.0(31.9-85.0) | 59.0(38.2-85.9) | 73.5(46.2-96.0) | 85.5(51.1-17.2) | 88.7(54.8-179.2) | 105.1(61.2-195.4) |
| > 75% to <100% wood use | 13 | 44.3(23.8-84.5) | 50.1(26.9-85.1) | 67.5(38.9-97.4) | 74.1(32.8-87.5) | 77.5(36.0-89.6) | 81.3(42.5-97.4) |
| 50% to 75% wood use | 21 | 52.1(36.2-93.6) | 54.9(39.8-99.1) | 76.6(43.5-116.8) | 92.2 (35.2 – 160.8) | 94.5 (37.6 – 163.9) | 119.0 (48.1-175.7) |
| 100% LPG use | 63 | 49.1(33.0-90.5) | 53.1(36.6-93.0) | 66.0(47.1-108.5) | 44.6(30.6-78.2) | 47.6(35.1-81.2) | 57.0(44.4-93.8) |
| p-value<sup>1</sup> |  | 0.441 | 0.700 | 0.700 | <0.007 | <0.007 | <0.054 |
| **Immediate Outdoors<sup>2</sup>** |  |  |  |  |  |  |  |
| 100% wood use | 29 | 55.8(44.3-73.3) | 57.1(44.5-75.8) | 61.3(47.7-78.4) | 114.9(81.8-247.5) | 115.1(83.4 - 247.9) | 115.7(87.0-254.7) |
| > 75% to <100% wood use | 8 | 42.5(24.6-106.2) | 45.4(25.7-107.5) | 57.5(32.0-111.20) | 83.4(39.1-175.9) | 84.6(43.0-176.5) | 87.2(52.8-177.6) |
| 50% to 75% wood use | 11 | 48.6(30.7-113.2) | 44.1(31.0-114.0)<br>11 | 59.1(34.4-116.8) | 57.7 (39.1 – 116.9) | 57.7 (39.1 – 116.9) | 66.2 (39.4 – 119.3) |
| 100% LPG use | 45 | 57.4 (34.9-111.2) | 58.1 (38.5-113.9) | 62.8(43.6-119.0) | 48.7(31.1-86.2) | 50.2(33.4-88.5) | 54.7(39.5-98.2) |
| p-value<sup>1</sup> |  | 1.0 | 1.0 | 1.0 | <0.001 | <0.001 | <0.001 |

<sup>1</sup> Significance of the comparison between 100% wood use and 100% LPG use using the Mann-Whitney U Statistic
<sup>2</sup> Monitors were not fixed in some of the kitchen monitoring session

PM_2.5_ concentrations in the kitchen and immediate outdoors of households using only wood for cooking with and without a chimney (when the kitchen is located in the house) are given in table 3. In households without a chimney (n = 8) using only wood (100%), the median PM_2.5_ concentration in the kitchens was more than three-fold higher than concentrations in households having a chimney (n = 21)(182.1 μg/m^3^ (IQR = 126.5 – 455.2μg/m^3^ vs. 182.1μg/ m^3^ (IQR = 126.5 – 455.2μg/m^3^)); the median outdoor PM_2.5_ concentration was two-fold higher in households without a chimney as compared to households with a chimney (245.9 μg/m^3^ (IQR = 72.5 – 641.7 μg/m^3^) vs. 105.7μg/m^3^ (IQR = 83.4– 195.8 μg/m^3^)).

**Table 3:** PM_2.5_ concentration in the kitchen and immediate outdoors of households cooking indoors and using only wood by presence of a chimney.

|  | Kitchen PM<sub>2.5</sub> concentration<br>Median (IQR) | Immediate Outdoors<br>Median (IQR) |
| --- | --- | --- |
| With chimney (n = 21) | 182.1 µg/m<sup>3</sup> (126.5–455.2<br>µg/m<sup>3</sup>) | 105.7 µg/m<sup>3</sup> (83.4– 195.8<br>µg/m<sup>3</sup>) |
| Without chimney (n = 8) | 635.4 µg/m<sup>3</sup> (473.7 –<br>5221.4µg/m<sup>3</sup>) | 245.9 µg/m<sup>3</sup> (72.5 – 641.7<br>µg/m<sup>3</sup>) |

A liner regression model was fitted to log transformed PM_2.5_ concentrations to assess important determinants of PM_2.5_ concentrations during cooking in houses where the kitchen was located in the main building of the house (Table 4). Fuel type and stove type, having a chimney in a fair or good condition, ratio of open space and total volume of kitchen, and baseline PM_2.5_ concentration were significant predictors of PM_2.5_ concentrations during cooking.

**Table 4:** Determinants of PM_2.5_ concentrations in Sri Lankan kitchens situated in the main house and using a single energy source (LPG or Wood))

| Variable | Regression coefficient | Standard error of regression coefficient | 95% confidence interval | Significance |
| --- | --- | --- | --- | --- |
| Baseline Kitchen PM<sub>2.5</sub><sup>1</sup> | 1.468 | 1.116 | 1.182-1.825 | 0.001 |
| Fuel type and stove type<sup>2</sup> | 3.394 | 1.149 | 2.577-4.469 | <0.001 |
| Any outdoor source (Burning, factory, road with in 50m etc. ) | 1.341 | 1.532 | 0.575-3.127 | 0.493 |
| Having a chimney and they are good or fair condition<sup>3</sup> | 1.889 | 1.233 | 1.247-2.860 | 0.003 |
| Ratio of open spaces in the kitchen and total volume in the kitchen<sup>4</sup> | 1.046 | 1.017 | 1.012-1.082 | 0.008 |
| Incense burning, mosquito coil or smoking<sup>5</sup> | 0.652 | 2.538 | 0.103-4.135 | 0.647 |
| (Constant) | 0.020 | 1.805 |  |  |

1. Log transformed data
2. Traditional wood stove and wood fuel use; improved wood stove and wood fuel, LPG stove and LPG fuel
3. Chimney condition is poor functionality; chimney present and they are in fair or good functionality
4. Ratio between space opens outdoors from the kitchen (height, width, and length of all the windows and doors open at the monitoring period) and total size of the kitchen (height, width, and length of the kitchen)
5. Indoor other sources

## Discussion

We compared PM fraction concentrations (PM_1,_ PM_2.5_ and PM_10_) in different microenvironments of Sri Lankan households using wood and LPG for cooking. PM concentrations at baseline and during cooking were extracted from the recordings of the air quality monitors. PM concentrations in the first 10 minutes of monitoring were used as the baseline PM concentration. Discarding the data in the window period between the end of baseline readings and the start of the cooking period would have ensured that changes in PM concentrations due to cleaning and setting up of the cook stove would not have been included. Obtaining a baseline value ensured that PM emitted from other sources including outdoor sources were adjusted for in the measurement. The cooking period in none of the households exceeded the monitoring period.

The majority of the households used more than one cooking fuel. Of the households that used a secondary cooking fuel, 44% and 11% of households used electricity and wood (up to 25% of cooking time), respectively. Results show that the households using wood 50% or more of the time but less than 100% of the time have lower PM_2.5_ concentrations in all household microenvironments as compared to households that use 100% wood for cooking. According to the energy ladder, electricity is considered to be the cleanest and most convenient type of energy, but expensive[7] Except one household, all the households in present study had electricity for lighting purpose. However only a single household predominantly used electricity (i.e., ≥ 75%) as a cooking fuel. This may due to cost as well as other factors such as relative availability of wood and cultural believes[2]. However, 43% and 12% of the households uses dry vegetation and wood respectively as the secondary cooking fuel. Some households uses more than one energy type as the secondary cooking fuel type as shown in the table 1.

As reported by other similar studies[10, 11], highest concentration of PM was reported from the wood using households. In fact, PM_2.5_ is several folds higher in wood using kitchens as compared to LPG using kitchens, and 24hour guideline value of World Health Organization (i.e., 25 μg/m^3^)[12].

Cooking with solid fuel increase the PM levels not only with in the household. It increases the immediate outdoor PM levels by more than two folds. Similarly, other studies prove the increase of outdoor air pollutant concentrations in solid fuel use[7].This confirms the solid fuel use finally contribute to the outdoor air pollution perhaps contributing to climate change. The black carbon, a component in the PM emit from solid fuel is considered to be the strongest short lived absorber of solar radiation in the atmosphere contributing to global warming seconding only to long lived greenhouse gases [13].

Present study shows that the PM_2.5_ pollution in the kitchen also increases the pollution in the other micro environment in the households. This shows, even if the household members do not come to the kitchen but stay in living room and sleeping room are exposed to high PM pollution.

Living room and sleeping room PM concentration during the cooking period is lower than the both kitchen and immediate outdoor. A similar study in Shanxi, China reported that the values in the rooms (114 ± 81μg/m^3^)are lower than the kitchen (376 ± 573μg/m^3^) but room values were higher than the outdoor values (64 ± 28μg/m^3^)[14]. This different may be due to structural differences of the household, climatic differences in two countries and differences in monitoring location.

Chimney is an important determinant of household air pollution specifically when the wood is used as the single cooking energy (Table 3). Present study shows PM_2.5_ concentration increases more than three folds in households not having a chimney and rely only on woodand cooking inside the main building irrespective other characteristics. A previous studies in Sri Lanka [5] and in other solid fuel using countries [15]show chimney use reduce the kitchen PM_2.5_ concentrations. Further, study reported comparatively higher immediate outdoor PM_2.5_concentration among the households not having a chimney as compared to households having a chimney (245.9 μg/m^3^ (IQR = 72.5 – 641.7 μg/m^3^Vs. 105.7μg/m^3^ (IQR = 83.4– 195.8 μg/m^3^). Although the results need to interpret cautiously, this may due to the fact that the chimneys emit the pollutant to the above the roofs of the households, where as kitchens not having the chimneys leaks the pollutants out of the doors and windows at the level of breathing zone (i.e., at the level of PM monitoring).

As a limitation, this study does not monitor the personnel PM exposure of household members thus challenging to quantify the personnel exposure levels. Nevertheless, study monitor the PM concentration in different microenvironments and immediate outdoor is a strength. In the practical context, considerable number of households used more than one cooking fuel. Although we have collected this information with the relative percentage of each fuel type, it may mislead the air quality levels reported from each energy type. As a remedial measure, we analyzed in depth only the households which uses only a single energy type for cooking during the monitoring session or categorized as households used > 75% to <100% wood use and 50% to 75% wood use.

## Conclusions

PMconcentrationsare several folds higher during the cooking with wood as compared to LPG in kitchens, living room and sleeping room. Immediate outdoor PM concentration is higher than the sleeping and living room PM concentrations. Among the households use wood and cook in the main building increases the PM_2.5_ concentration by more than three folds during the cooking periods. Households not having a chimney had higher PM_2.5_ concentrations at immediate outside of the household as compared to households having a chimney. The important factors that determined the PM_2.5_ concentrations in cooking indoors were 1) Fuel type and stove type (2) availability of a chimney and their functional status (3) Ratio between open space and total space of kitchen (4) PM_2.5_ concentration at the non-cooking time.

